# Evaluating Sequence and Structural Similarity Metrics for Predicting Shared Paralog Functions

**DOI:** 10.1101/2024.10.11.617835

**Authors:** Olivier Dennler, Colm J. Ryan

## Abstract

Gene duplication is the primary source of new genes, resulting in most genes having identifiable paralogs. Over evolutionary time scales, paralog pairs may diverge in some respects but many retain the ability to perform the same functional role. Protein sequence identity is often used as a proxy for functional similarity and can predict shared functions between paralogs as revealed by synthetic lethal experiments. However, the advent of alternative protein representations, including embeddings from protein language models (PLMs) and predicted structures from AlphaFold, raises the possibility that alternative similarity metrics could better capture functional similarity between paralogs. Here, using two species (budding yeast and human) and two different definitions of shared functionality (shared protein-protein interactions, synthetic lethality) we evaluated a variety of alternative similarity metrics. For some tasks, predicted structural similarity or PLM embedding similarity outperform sequence identity, but more importantly these similarity metrics are not redundant with sequence identity, i.e. combining them with sequence identity leads to improved predictions of shared functionality. By adding contextual features, representing similarity to homologous proteins within and across species, we can significantly enhance our predictions of shared paralog functionality. Overall, our results suggest that alternative similarity metrics capture complementary aspects of functional similarity beyond sequence identity alone.

**GRAPHICAL ABSTRACT:** 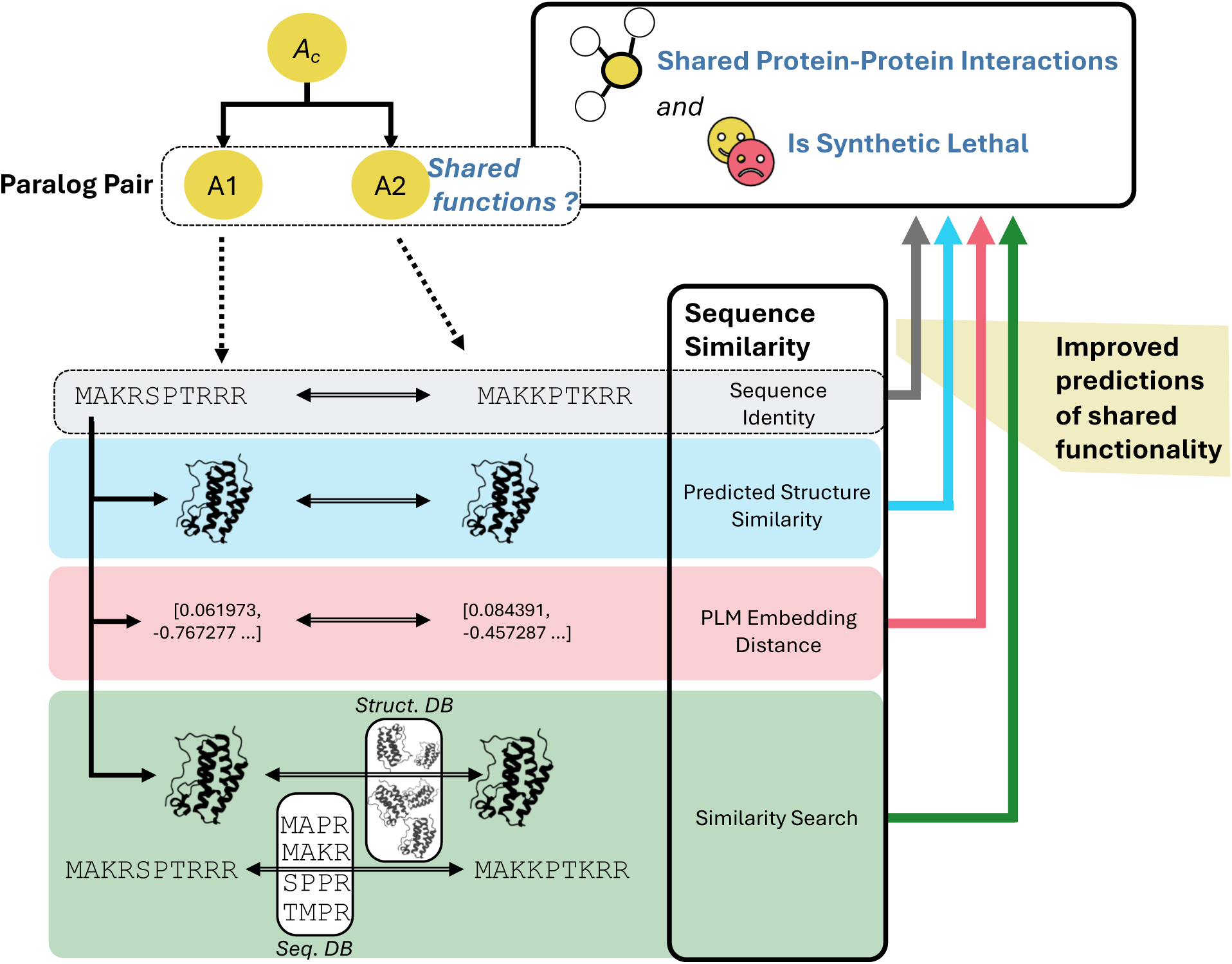

## INTRODUCTION

Comparisons of protein sequences provide the foundation for much of modern biology, allowing us to infer evolutionary relationships between genes and predict functions for homologous proteins (1–6). Such sequence comparisons are not just made across species, for inferring functions of orthologs, but also within species, for paralogs (genes that arose from the duplication of a common ancestral gene). Most genes, including approximately 70% of human genes, have identifiable paralogs and therefore understanding the shared and divergent functions of paralog pairs is crucial to understanding gene function and the genotype-phenotype map (7).

Over evolutionary time scales, pairs of paralogs accumulate mutations, leading to divergence in their protein sequences and potentially in their function. However, even after diverging for hundreds of millions of years, many pairs of paralogs still perform similar functions in the cell and in some cases exhibit sufficient functional similarity that they can phenotypically compensate for each other’s loss (8–10). Such compensatory relationships can be revealed via perturbation experiments – often it is possible to mutate each paralog individually with little fitness consequence but their mutation in combination results in cell death, a phenomenon known as *synthetic lethality*. These compensatory relationships between paralogs contribute to *genetic robustness* – the ability of cells and organisms to withstand genetic perturbation (11, 12) – and can potentially be exploited for the development of targeted therapies in cancer where deletion/mutation of one paralog may render tumour cells vulnerable to inhibition of another (8, 13–22).

Previous work has demonstrated that the amino acid sequence identity between pairs of paralogs is predictive of synthetic lethality, suggesting that sequence similarity is a reasonable proxy for the functional similarity of paralogs (15, 20, 23). Recent computational advances have led to the development of new protein sequence representations, such as AlphaFold-predicted protein structures (24, 25) and Protein Language Model (PLM) embeddings (26, 27). These representations, which are essentially derived from protein sequences, are now being widely used to predict functions from protein sequences (28–30). In this work, we assess whether similarity metrics derived from these alternative protein representations can be used to predict whether two paralogs share functions.

We model this as a binary classification problem – can similarity metrics derived from sequence, structure, and PLM embeddings effectively identify pairs of paralogs where both paralogs share functions (Figure 1)? We use two definitions of *shared functions* between paralog pairs – shared protein-protein interactions (PPIs), indicating that the two paralogs occupy similar spaces in the protein-protein interaction network, and synthetic lethality (SL), indicating that the paralogs can phenotypically compensate for each other’s loss (31). By focusing on these two specific definitions, we aim to better understand how various sequence similarity metrics correlate with functional conservation in paralogs, and to provide insights into the broader question of sequence-function relationships. To ensure generalisability of our findings, we perform our evaluation using paralogs from two species – humans and the budding yeast *Saccharomyces cerevisiae*, both of which have comprehensive protein-protein interaction maps as well as systematically generated synthetic lethal interaction data available.

**Figure 1.**
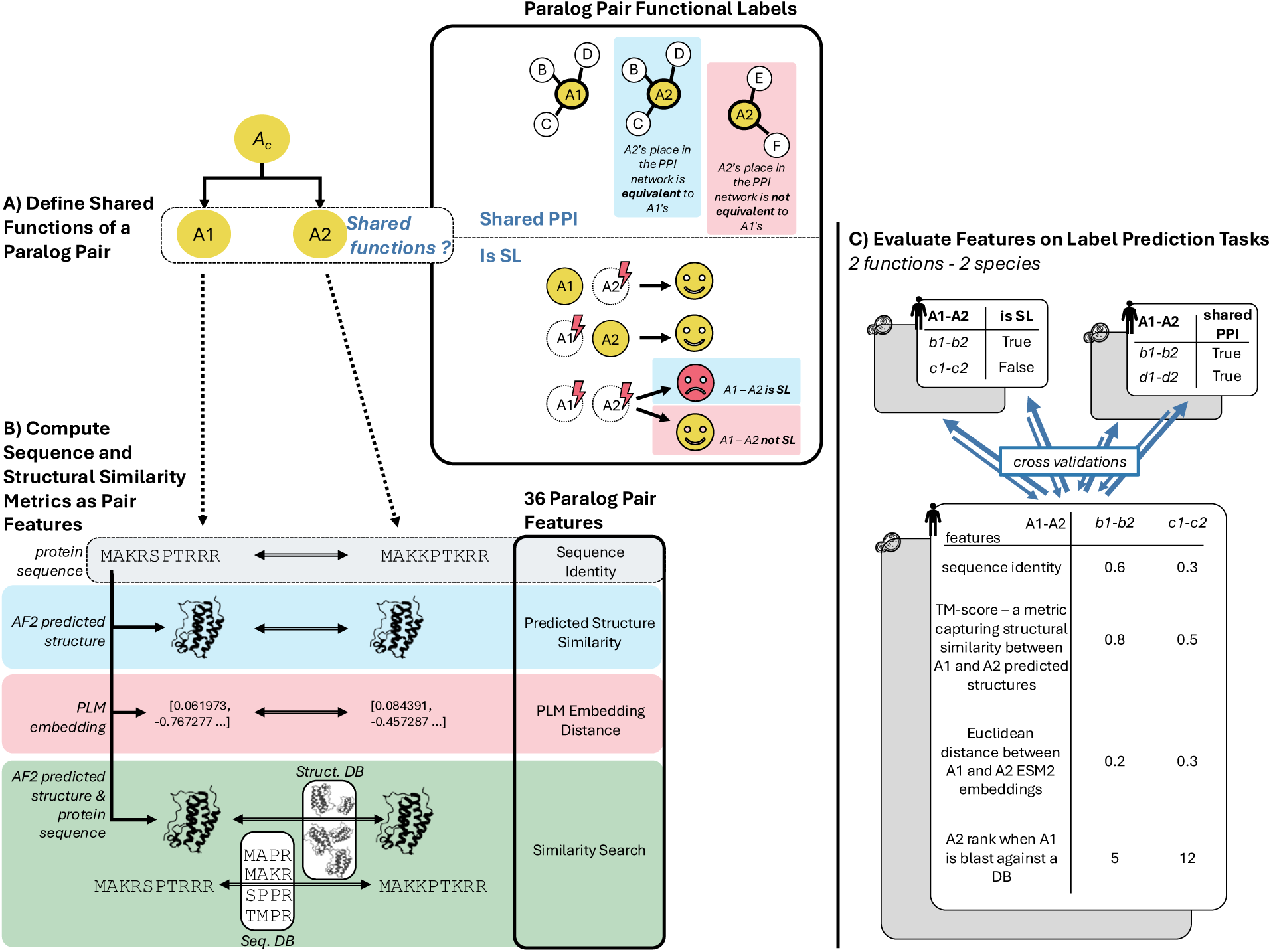
Experimental overview – evaluating sequence and structural similarity metrics for predicting shared paralog functions. **A)** A paralog pair A1-A2 is annotated based on their ability to perform shared functions. Two distinct labels are used: shared PPI (the two genes of the pair share a significative overlap of their PPI partners), and synthetic lethality (SL, simultaneous loss of both genes of the pair induces death cell when their individual loss has no effect). **B)** For every pair of paralogs we include amino acid sequence identity as a baseline and calculate features from comparison of their predicted structures (such as the TM-score) and their Protein Language Model (PLM) embeddings (such as cosine similarity). We also calculate features derived from searches of each paralog against multi-species databases of protein sequences and predicted protein structures. These features allow us to evaluate how similar the paralogs are to each other with respect to other proteins in the same species (paralogs) and across other species (orthologs) (e.g. is A2 the most similar protein in the database to A1 or are there 10 more similar paralogs?). **C)** Using a cross-validation approach we evaluate the ability of each individual feature to predict two definitions of *shared functions* (PPI/SL) across two species (yeast and human). Using machine learning we also evaluate the predictive power of feature subsets (e.g. all structural features) and all features combined.

Our evaluation across four datasets (2 definitions x 2 species) reveal that sequence similarity features derived from alternative protein representations provide non-redundant but complementary functional information that is not captured by sequence identity alone. While individual similarity measures perform better in specific scenarios, the combination of all features in machine learning classifiers consistently offers a more accurate prediction of whether two paralogs share functions. We further show that features derived by searching for homologous proteins within and across species provides additional information beyond simple pairwise comparison and can aid the identification of paralog pairs where both paralogs share functions.

## MATERIAL AND METHODS

### Building four paralog pair classification datasets

#### Human and yeast paralog pairs lists

We obtained 124,813 human (*Homo sapiens*; genome assembly GRCh38.p14) and 5,673 yeast (*Saccharomyces cerevisiae S288c*; genome assembly R64-1-1) protein-coding paralog pairs and their corresponding amino acid sequence identities from Ensembl release 111 (32). Each gene was mapped to its corresponding UniProt entry. Paralog pairs for which all AlphaFold and Protein Language Model features could not be computed were subsequently filtered out, resulting in a final total of 107,103 paralog pairs for humans and 5,673 for yeast.

#### Annotating paralog pairs as sharing functions based on protein-protein interaction comparison

For the shared PPI labels, we assessed whether paralog pairs share functions based on the overlap of their protein-protein interactions (PPI). We used systematically generated PPI networks for this. For humans, PPI data were obtained from BioPlex 3.0 (HEK293T cell AP-MS) (33), including only interactions where the paralog served as the bait. For yeast, PPI data were sourced from the Michaelis et al. (34) yeast interactome. We measured the overlap of interactors for each paralog pair (A1-A2) using the −log10 p-value from a Fisher’s exact test (FET) to determine the significance of the overlap. The FET was conducted by comparing the observed number of shared protein-protein interactions (PPIs) between two paralogs to the total set of all detected proteins involved in interactions within the experiment, serving as the background population. Paralog pairs were classified as “sharing functions” if the −log10 p-value from the FET was greater than or equal to −log10(0.05/number of pairs in the species), indicating significant overlap. Conversely, pairs were classified as “not sharing functions” if the −log10 p-value from the FET was less than or equal to −log10(0.05), indicating non-significant overlap. Pairs with nominally significant overlap that did not meet significance criteria after Bonferroni correction for multiple hypothesis correction were considered ambiguous and removed from the datasets.

#### Annotating paralog pairs as sharing functions based on synthetic lethality

In humans, SL labels were assigned using the method described in De Kegel *et al* (35) (training dataset). Briefly, synthetic lethality between paralogs was called by analysing CRISPR screens in a panel of cancer cell lines and using a linear regression model to identify associations between loss (via deletion, silencing, mutation) of one paralog and sensitivity to the inhibition of its counterpart. De Kegel *et al* (35) only evaluated paralog pairs from smaller families (<20 paralogs) and that met a minimum sequence identity threshold (20%). Here we used the same pipeline but assessed all paralog pairs in Ensembl 111 without applying sequence identity or family size filters. In yeast, SL labels were assigned based on the SGA (Synthetic Genetic Array) genetic interaction dataset (36), which involves pairwise screening of non-essential genes (NxN). Pairs with a genetic interaction score ≤ −0.08 and a p-value ≤ 0.05 were classified as SL, while the remaining screened pairs were classified as non-SL. Non-screened pairs were systematically filtered out.

### Computing paralog pair features

#### Features derived from comparing predicted structures

For each paralog pair (A1-A2), we retrieved the AlphaFold2 predicted structures for A1 and A2 canonical isoforms from the AlphaFold Database (24, 37) (UP000005640_9606_HUMAN_v4 for human; UP000002311_559292_YEAST_v4 for yeast). We then used Foldseek (38) (version 6.29e255) to search the structure of A1 against the entire AlphaFold Database. The resulting Foldseek ranked search output is similar to a BLAST (1) output table, as it contains alignment results between a query protein and potential target proteins, ranking them based on alignment scores. For A1 as the query, we extracted the alignment scores corresponding to A2 as the target. This process was repeated with A2 as the query and A1 as the target. The following pair features were defined based on the alignment scores: fident, TM-score(s), LDDT, bits, e-value, probability, and alignment length. For each feature, the minimum value between the two alignments (A1 as query and A2 as query) was used. In the minority of cases (3.4%) where neither A1 nor A2 provided an alignment with their respective paralog, the pairs were systematically filtered out.

#### Features derived from database searches

The Foldseek ranked search output, previously used for assessing pairwise structural similarity for all paralog pairs, was also utilized to define structural similarity search features. For sequence similarity search features, we employed MMseqs2 (39) (version 13.45111) in a similar manner to Foldseek. We chose MMseqs2 due to its similarity in implementation to Foldseek, which helps minimize algorithmic differences and ensures fair comparisons. We searched the A1 UniProt sequence against the UniProtKB/Swiss-Prot (40) (release 2023_04 of 13-Sep-2023) sequence database and identified A2 in the resulting ranked search table. We used the similarity search results to define the following three features: the rank of A2 in the ranked search results, the number of proteins from the same species as A1-A2 that rank higher than A2 (paralogs), and the number of species that possess proteins ranking higher than A2 (species). This process was repeated with A2 as the query and A1 as the target, and the final pair features were determined by taking the minimum values observed between the two alignments (A1 as query and A2 as query). In the minority of cases (3.4%) where neither A1 nor A2 provided an alignment with their respective paralog, the pairs were systematically filtered out.

#### Distances between Protein Language Model embeddings as pair features

We used two sources of PLM embeddings – ProtT5 and ESM2. For ProtT5 (27) (ProtT5-XL-U50 model), “per-protein” pre-encoded embeddings for A1 and A2 were retrieved from the UniProt database and compared using the following distance metrics: cosine, Euclidean, Manhattan, and TS-SS (Similarity-Sector Similarity) (41). For ESM2 (42), the UniProt protein sequences of A1 and A2 were encoded using the “esm2_t48_15B_UR50D” model. Since the resulting embeddings were of different, incomparable sizes, we generated “fixed-size” protein embeddings using four methods based on the implementation by Yeung et al. (43): beginning-of-sequence tokens, end-of-sequence special token, mean of both special tokens, and mean of all residue tokens. These fixed-size embeddings were then compared using cosine, Euclidean, Manhattan, and TS-SS distances.

### Evaluating pair features on label predictions

#### Pearson’s correlation coefficients

To evaluate the relationships between the different features, Pearson correlation coefficients were calculated using Python 3.11.4 and the pandas 2.1.3 package. Since the features could be either positively or negatively correlated, we presented the absolute values of the correlation coefficients for clarity.

#### Evaluating individual features for predicting labels

We evaluated each feature across all four datasets for their ability to predict whether paralog pairs share functions. Using 4-fold cross-validation, we first standardized the feature values using z-score normalization (sklearn version 1.3.2 used for all machine learning analyses). Then, we assessed the performance of simple logistic regression classifiers (max iteration=1000), each containing only one feature at a time, to predict the paralog pair labels. The performance was measured by computing the area under the receiver operating curve (AUROC) for each feature, and we presented both the individual AUC values and the mean AUC across the folds.

#### XGBoost classifiers evaluation on label predictions

To evaluate the predictive power of each *set* of features(predicted structure similarity, PLM embeddings, similarity search), rather than individual features, we used XGBoost classifiers. As before, we applied 4-fold cross-validation and standardized all feature values. We assessed the performance of XGBoost classifiers (n_estimators=600, random_state=8, learning_rate=0.1, colsample_bytree=0.5, use_label_encoder=False, eval_metric=’logloss’, xgboost version 2.0.3) that incorporated various subsets of features to predict the paralog pair labels. The performance of these classifiers was measured by computing their area under the receiver operating curve (AUROC). We presented the mean AUROC across the folds, along with the receiver operating curve and the precision-recall curve for each set of features.

### Constructing additional datasets for Gene Ontology-based paralog pair classification

#### Gene ontology terms retrieval

Human Gene Ontology (GO) terms were sourced from UniProt (reference proteome data, 2023-07-12) (40). For yeast, GO terms were retrieved from the Saccharomyces Genome Database (SGD, 2024-03-25) (44). Only experimental GO terms were retained, filtered by evidence codes: EXP, IDA, IEP, IGI, IMP, IPI, TAS, and IC. Paralog genes were then annotated with their corresponding experimental GO terms.

#### GO semantic distances between paralogs

The GOGO software (45) was used to calculate semantic distances between the two genes of each paralog pair, generating separate distances for Biological Processes (BPO), Molecular Functions (MFO), and Cellular Components (CCO).

#### Paralog pairs classification based on GO semantic distances

Paralog pairs were classified using fixed thresholds: pairs with semantic similarities ≥ 0.9 were labelled as sharing functions, while those with similarities ≤ 0.3 were labelled as not sharing functions. Pairs with semantic similarities not meeting the above mentioned criteria were excluded from the analysis. This process resulted in six datasets: three for human (BPO, MFO, CCO) and three for yeast (BPO, MFO, CCO), where paralog pairs were classified based on their respective semantic similarity.

## RESULTS

### Defining paralog pairs with shared functions using protein-protein interaction networks and synthetic lethality

We first obtained a list of 107,103 paralog pairs in humans (*Homo sapiens*) and 5,673 paralog pairs in budding yeast (*Saccharomyces cerevisiae)* from Ensembl111 (32). We then annotated each set of paralog pairs according to whether or not they share functions. We reasoned that pairs of paralogs could be viewed as having shared functions if they 1) could interact with the same protein-protein interaction (PPI) partners or 2) could phenotypically compensate for each other’s loss, as evidenced by synthetic lethality (Figure 1A). For the PPI features, we made use of systematically generated PPI networks (Bioplex for humans ; social interactome for yeast) (33, 34) to avoid the ascertainment bias associated with networks that aggregate interactions from multiple sources (46). Using these networks, paralog pairs were annotated as having shared functions if they shared a significant fraction of their PPI partners (see Methods), resulting in 3.6% of pairs in humans and 10% of pairs in yeast labelled as having shared functions (Figure 2A). For SL pairs, we made use of different sources of labels for each species. In budding yeast, a near complete map of all genetic interactions between pairs of protein-coding genes has been generated (36) which includes the vast majority of paralog pairs. We used this to annotate 10.9% of yeast paralog pairs as having shared functions (Figure 2A). No similar comprehensive genetic interaction map exists for humans, but genome-wide CRISPR screens in genetically diverse cancer cell lines can be used to identify robust and unbiased synthetic lethal and non-synthetic lethal pairs (35). Using this approach, we annotated 0.5% of human paralog pairs as synthetic lethal (Figure 2A). We note that in each dataset a subset of gene pairs was left unannotated due either to lack of data (e.g. missing CRISPR gene scores for some genes) or ambiguous calls (e.g. a nominally significant overlap in PPI partners that did not meet the threshold for significance, see Methods).

**Figure 2.**
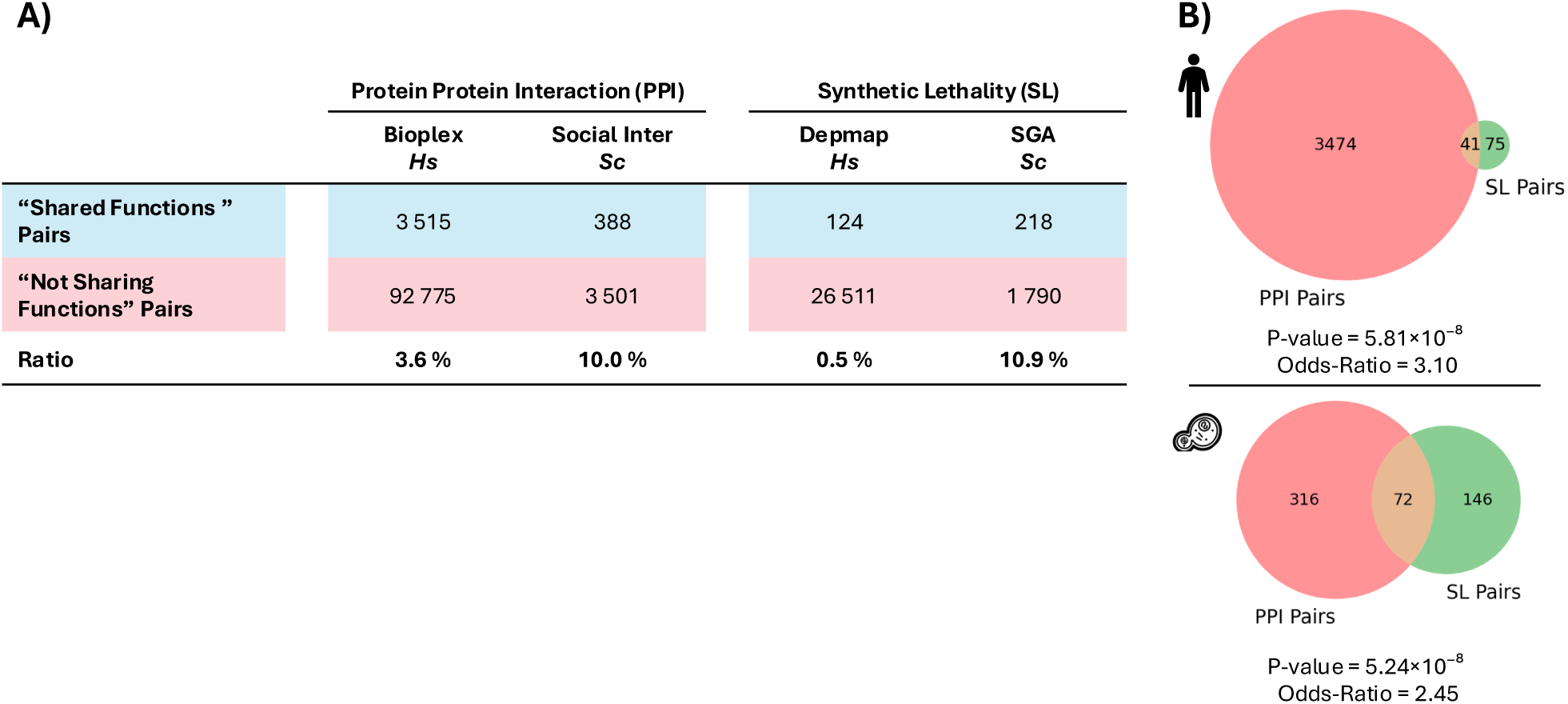
Details of “shared functions” labels in two species. **A)** Table showing the counts of paralog pairs labelled according to four “shared functions” class annotations (one PPI and one SL dataset for each species, humans (*Hs* – *Homo sapiens*) and budding yeast (*Sc* – *Saccharomyces cerevisiae*)). **B)** Venn diagram illustrating the overlap between pairs labelled as sharing functions based on PPI and SL in humans and in budding yeast, highlighting the small but significant level of agreement and the varying scales between the different datasets. P-value and Odds-ratio from a Fisher’s Exact Test (FET).

Figure 2B shows the composition of the resulting four datasets. Although these datasets derive from the same lists of Ensembl paralog pairs, the labels within each species are defined using different functional criteria (PPI and SL) and so they are only partially overlapping. A pair labelled as “sharing functions” in one dataset might be labelled as “not sharing functions” or not labelled at all in another. We assessed the agreement between the two functional definitions using Fisher’s exact tests, revealing statistically significant relationships between the datasets of the same species (Figure 2B, PPI-SL p-value = 5.81×10⁻⁸ in humans ; PPI-SL p-value = 5.24×10⁻⁸ in yeast). These results suggest that different functional definitions based on distinct methodological approaches successfully capture common biological insights regarding paralog function sharing.

To evaluate various sequence similarity metrics for predicting paralogs sharing functions, we analysed each of these four paralog datasets separately. We aimed to assess the suitability of these metrics for specific annotations and to understand how different features perform across various functional annotations, species, dataset sizes, and label distributions.

### Evaluation of structural similarity between predicted structures to predict shared functions in paralogous genes

Protein sequences fold into 3D structures, enabling them to interact with their environment and perform their functions. Thanks to AlphaFold2 (24, 37) predicted protein structures derived from sequences are now widely available for many species including both yeast and humans. To assess whether structural similarity can predict if two paralogs perform the same function, we compared the AlphaFold2 predicted structures of paralog pairs. We used Foldseek (38), a recently developed tool designed for fast and efficient protein structure alignments, to align the AlphaFold predicted structures.

Foldseek is akin to BLAST but for protein structures. It enables fast structure alignments and quick searches for structural similarities across large protein databases. We evaluated the scores computed by Foldseek as features that define the pairwise structural similarity of the paralogs. We defined nine pairwise structural similarity features (Figure 3A, left as follows): amino acid sequence identity after Foldseek structural alignment (fident), three Template Modelling (TM) scores that measure global structural similarity (TM-score of the alignment; TM-score normalised by the query length; TM-score normalised by the target length) (47), Local Distance Difference Test (LDDT) (48), which measures local structure similarity, and four additional scores from the Foldseek alignment (e-value, Bits score, Probability, and alignment length).

**Figure 3.**
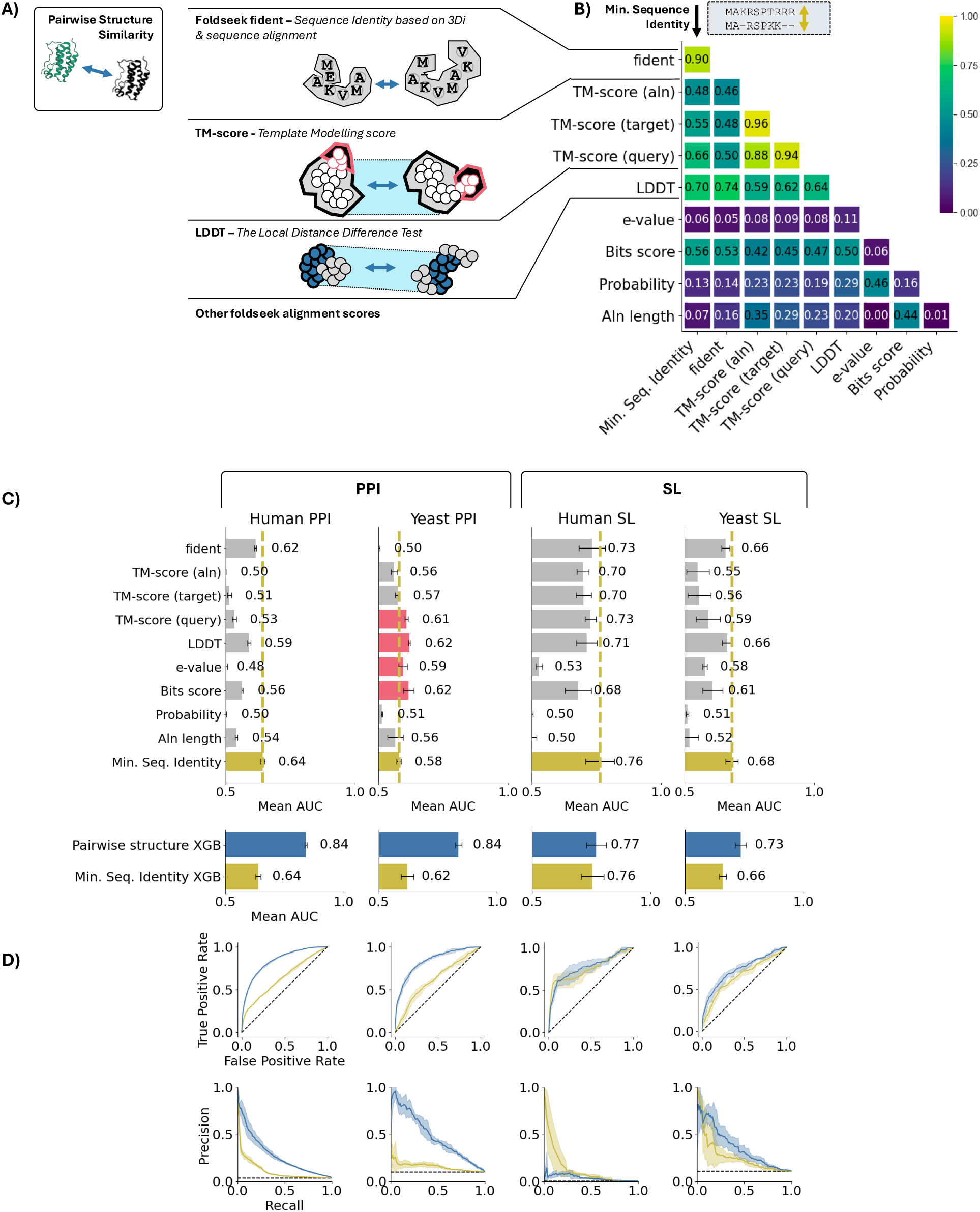
Pairwise structural comparison features provide non-redundant information about shared paralog function. **A)** Overview of nine predicted structural similarity features derived from Foldseek alignment between AlphaFold predicted structures of the two paralogs. **B)** Pearson correlation coefficients (absolute values) between these nine features and sequence identity in the 100k human paralog pairs from Ensembl 111. **C)** Mean AUC values comparing the nine individual structural features with minimum sequence identity (in yellow) for predicting shared functions across the four datasets using four fold cross validation. Features with mean AUC greater than the mean AUC of sequence identity are shown in red. Error bars show the standard deviation of AUC values from cross-validation. **D)** Performances of an XGBoost classifier using the nine predicted structure similarity features plus sequence identity (in blue) compared to a classifier using solely sequence identity (in yellow). Top chart shows the mean AUC (AUROC) on each dataset, bottom shows ROC and precision-recall curves for each datasets.

Considering these nine pairwise features are derived from AlphaFold predicted structures, and hence essentially from protein sequences, we investigated whether they contain redundant information already captured by sequence identity or if they provide novel information. We first computed Pearson correlation coefficients between the selected features for all human paralog pairs (Figure 3B). We found that only fident displayed a high correlation (Pearson correlation of 0.9) with standard sequence identity, while all other metrics were considerably lower. The high correlation with fident is expected, as it is effectively an amino-acid sequence identity measurement calculated from a pairwise alignment derived from aligned structures. Other structural similarity metrics displayed significantly lower correlation with sequence identity, e.g. LDDT score (0.7) and TM-score aln (0.48), suggesting that they capture non-redundant information. Similar observations were made for the correlations between calculated features in the yeast dataset as shown in Supplementary Fig. 1.

As these nine pairwise features capture pairwise structural divergence, they may also capture functional divergence. To evaluate their ability to predict whether two paralogs share functions, we performed a 4-fold cross-validation on each of our four datasets individually. This approach ensured that we assessed the predictive performance separately within each dataset. As shown in Figure 3C, most features had at least some predictive power (mean AUC > 0.5), though their performance varied across datasets. With the exception of predicting shared PPI in yeast, no structural similarity feature outperformed sequence identity at predicting shared functions. Some features (e.g. fident and LDDT) performed similarly or marginally worse than sequence identity at predicting SL in yeast and humans, but did not outperform it. For yeast PPIs, most structural features, led by LDDT and Bits score, outperformed sequence identity, which performed particularly poorly at this task (AUC = 0.58). This suggests that structural comparisons may be particularly valuable when sequence-based comparisons perform poorly such as an in the yeast shared PPI challenge. However, they might be unnecessary when sequence identity alone is sufficient, as seen in human SL.

Since the different structural features seem to capture various aspects of functional divergence, we developed an XGBoost classifier combining all nine predicted structure features with sequence identity. We evaluated this classifier’s performance using cross-validation on the four datasets and compared it to a classifier that relied solely on sequence identity. As shown in Figure 3D, this structure-based classifier outperformed sequence identity in most datasets, except in human-SL, where sequence identity alone performed exceptionally well. These results suggest that sequence identity and predicted structural similarities provide non-redundant information about shared paralog functions.

### Evaluation of distances between protein language model embeddings to predict shared functions in paralogous genes

Protein Language Models (PLMs) represent another AI application in biology, designed to learn the language of protein sequences from vast sequence datasets. They represent protein sequences as high-dimensional vectors (embeddings) in this learned language space. To evaluate the ability of distances between PLM embeddings to predict if two paralogs perform the same function, we investigated two commonly used Protein Language Models: ESM2 (42) (15B parameters ; trained on UniRef50) and ProtT5 (27) (a transformer using 3B parameters ; trained on UniRef50), along with various metrics for comparing the produced PLM embeddings.

ESM2 produces “per residue” embeddings that are not directly comparable at the protein level because their lengths vary based on the protein sequence. To make them comparable, we used four methods to compute fixed-size embeddings: beginning of sequence, end of sequence, mean of residue tokens, and mean of special tokens. For ProtT5, we directly obtained the fixed-size “per protein” embeddings available on UniProt. For each fixed-size embedding (five in total — four from ESM2 and one from ProtT5), we computed four different distances (cosine, Euclidean, Manhattan, TS-SS) between the embeddings of each paralog pair, resulting in a total of 20 PLM embedding distances used as pair features.

Given that these 20 features are derived from protein sequences, we investigated whether PLM embedding distances contain redundant information already captured by sequence identity. We first computed Pearson correlation coefficients between the selected features for all human paralog pairs (Figure 4B). While all feature pairs showed some correlation, only those corresponding to different distance metrics between the same fixed-size embeddings displayed high correlation (Pearson correlation > 0.8). In general, the correlation with protein sequence identity was only moderate (average Pearson correlation across all metrics being 0.51). The observed correlations suggest that the 20 PLM features capture non-redundant information, influenced by the choice of PLM, the method used to compute fixed-size embeddings, and to a lesser extent, the distance metrics used (similar observations were made in yeast, as shown in Supplementary Fig. 1).

**Figure 4.**
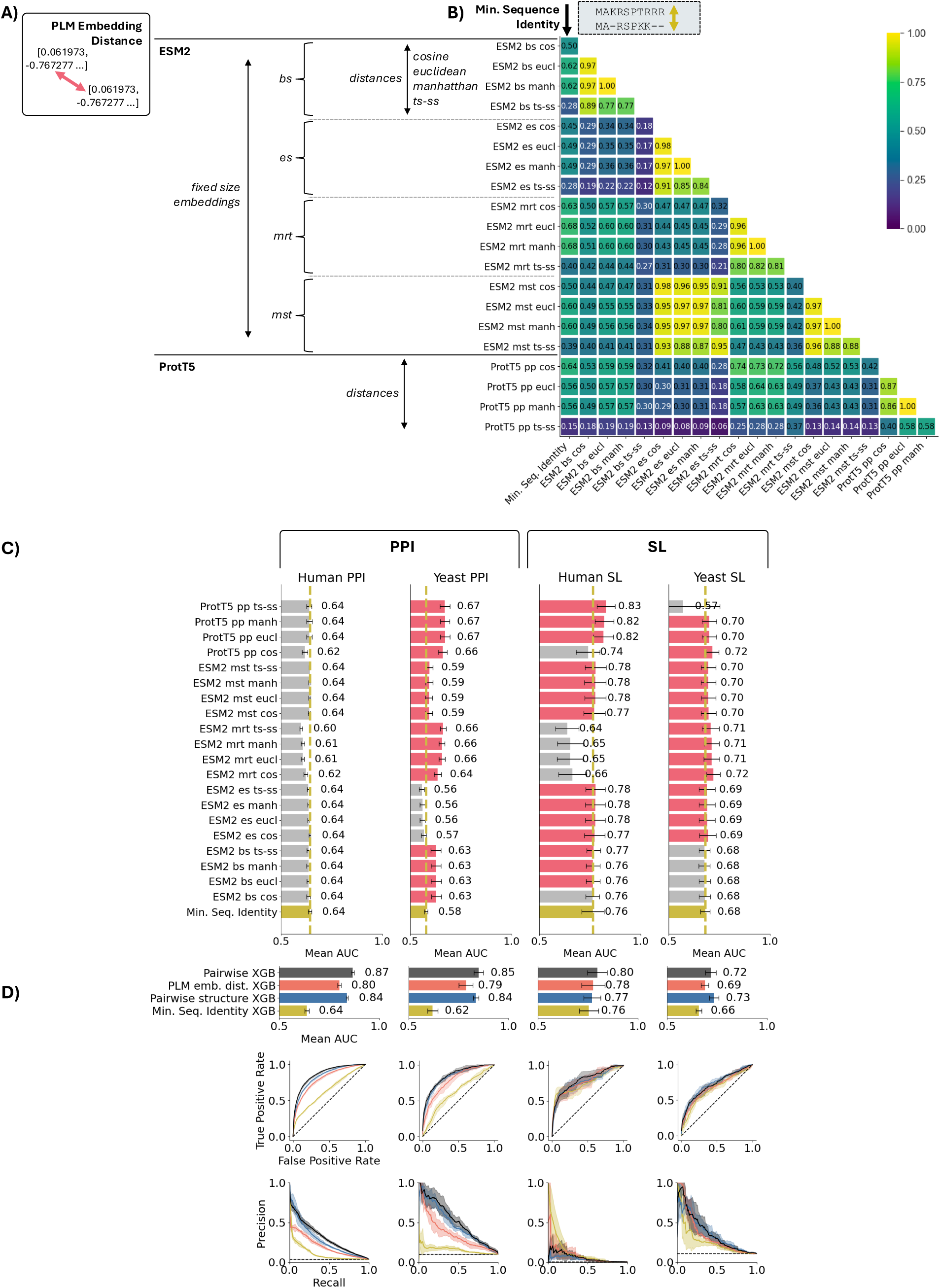
Distances between protein language model embeddings provide novel information about the functional similarity of paralog proteins. **A)** Overview of the twenty PLM embedding features derived from ESM2 and ProtT5 embeddings. ProtT5 embeddings are available directly in fixed sizes per protein, while ESM2 embeddings are outputted in varying sizes based on the initial protein sequence length. ESM2 embeddings are converted to fixed sizes using four different methods: bs: fixed size using the beginning of sequence token, es: fixed size using the end of sequence token, mrt: fixed size using the mean of residue tokens, mst: fixed size using the mean of special residue tokens. Four different distance metrics (cosine similarity, Euclidean, Manhattan, TS-SS) are used to compare each paralog pair’s embeddings for a given fixed size, resulting in 20 features. **B)** Pearson correlation coefficients (absolute values) between these 20 features and sequence identity in 100k human paralog pairs from Ensembl 111 revealing that the method of fixing the size of the embedding matters more than the distance metric. **C)** Mean AUC of the twenty individual features compared to minimum sequence identity (in yellow) for predicting shared functions across the four datasets using four fold cross validation. Features with mean AUC greater than the mean AUC of sequence identity are shown in red. Error bars show the standard deviation of AUC values from cross-validation. **D)** Performances of an XGBoost classifier using the 20 PLM features plus sequence identity (in dark orange) compared to a classifier using solely sequence identity (in yellow), the predicted structure similarity classifier (in blue), and a classifier with all 29 features combined with sequence identity (in grey) for 4-fold cross-validation on shared functions prediction across the four datasets. Top chart shows the mean AUC (AUROC) on each dataset, bottom shows ROC and precision-recall curves for each datasets.

As these 20 PLM pair features directly capture pairwise sequence similarity, we hypothesize they may also capture functional similarity. To evaluate their ability to predict if two paralogs share functions, we performed 4-fold cross-validations on the four datasets as we did for structural similarity metrics. As shown in Figure 4C, most PLM features had similar predictive power and performed comparably to each other across all four datasets. The biggest differences were observed between ESM2 and ProtT5 derived distances, with ProtT5 generally performing better, and also between the different ESM2 fixed-size embeddings, indicating that the method used to make the embeddings fixed-size matters more than the distance metrics used.

Since the different PLM features seem to capture functional similarity without being highly correlated, we combined all 20 PLM features with minimum sequence identity in an XGBoost classifier and evaluated its performance using cross-validation on four datasets. As shown in Figure 4D, this PLM classifier outperformed sequence identity alone in all datasets except human-SL, where it achieved equivalent performance.

To determine if the nine structural features and the 20 PLM features contain redundant or unique information, we built a third XGBoost classifier incorporating all previous 29 pairwise features. This classifier significantly outperformed sequence identity alone (Figure 4D), as well as the individual PLM and structural classifiers, across all four datasets. Combining these features resulted in substantial gains in predictive power for all classification challenges. These results suggest that sequence identity, PLM embedding distances, and predicted structural similarities all contain non-redundant functional information.

### Sequence - structure similarity searches against databases further capture novel information on paralog pairs sharing functions

Previous work has established that the likelihood of two paralogs displaying synthetic lethality is influenced not only by the similarity of the two proteins, but also by the number of other paralogs in the same family and whether or not the pair in question are the most similar within a larger family (35). The existence of other members within a larger family may also influence the likelihood of two proteins sharing protein-protein interactions. We hypothesise that within a given paralog family, if shared functions exist, the closer paralogs— those that are more similar—will be better candidates for sharing functions, while the distant ones—those that are less similar—will be less likely to share functions. Here, we aim to evaluate whether the presence of more similar sequences or structures in databases can predict if two paralogs share functions.

To determine the existence of more similar protein sequences, we first performed similarity searches using sequences and structural information. The two paralog sequences of a pair were queried against the UniProt database (40) using MMseqs2 (39). The resulting position of the paralog in the ranked search results was translated into different features: the number of more similar protein sequences among all homologs (rank), the number of more similar proteins within the same species (paralogs), and the number of species with more similar orthologs (species). Similarly, to identify more similar protein structures, we used Foldseek (38) to query the predicted structures of the paralogs against the AlphaFold database (24, 25, 37), resulting in three analogous features for structures. This approach yielded a total of six features capturing the existence of more similar sequences and structures (Figure 5A).

**Figure 5.**
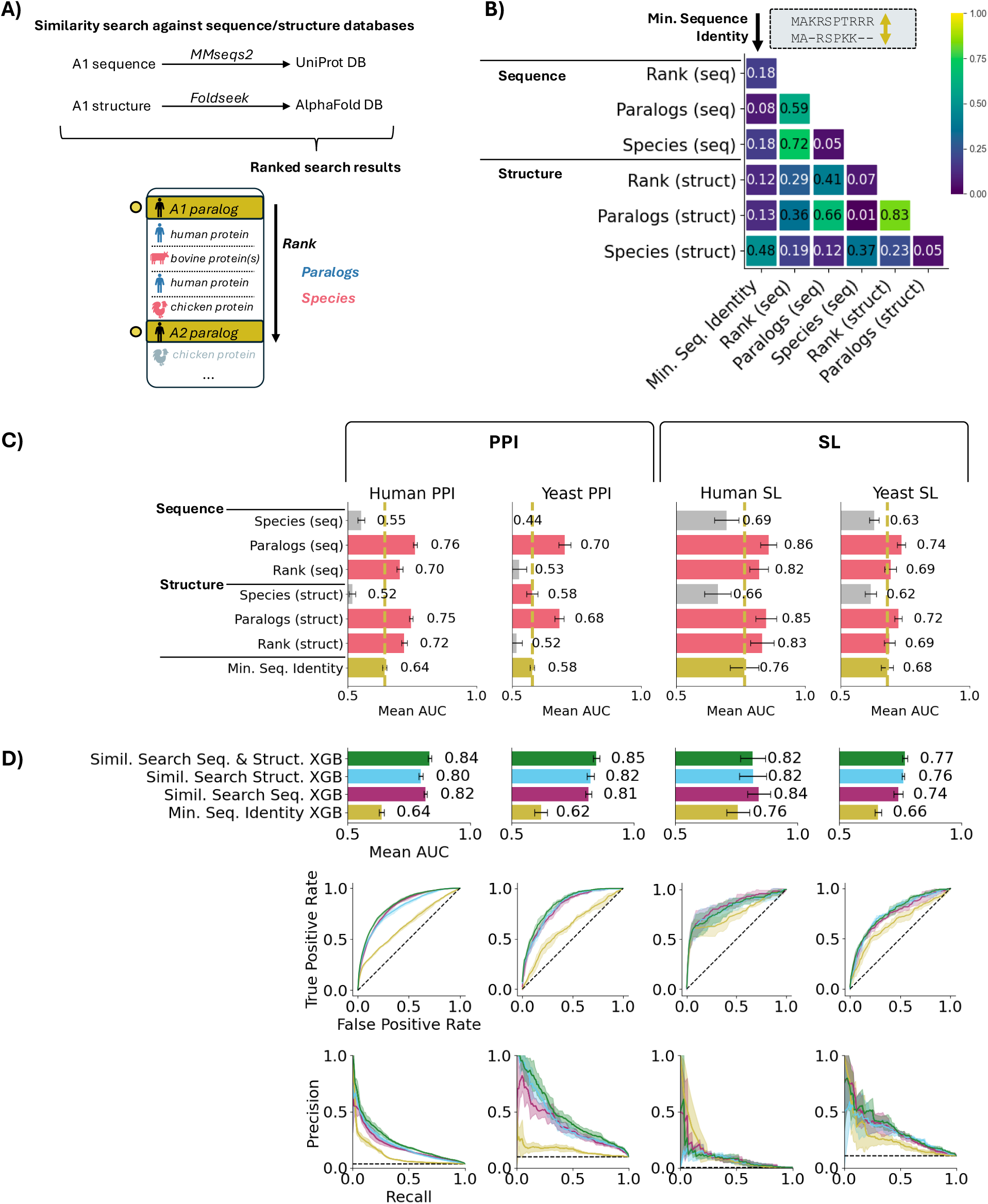
Sequence and structure similarity searches provide non-redundant information about shared paralog function. **A)** Overview of the similarity search features. We use MMseqs2 to search paralog sequences against the UniProt sequence database and Foldseek to search AlphaFold predicted structures of paralogs against the AlphaFold database. Features are extracted from the ranked search results, specifically focusing on the paralog rank (rank of A2 when A1 is the query). To capture more precisely the divergence within and across species captured by the database searches, we derive two additional features from the ranked search result table: “paralogs”: the number of proteins within the same species that rank better than A2, and “species”: the number of species possessing proteins (orthologs) that rank better than A2. These three feature pairs (rank, paralogs, species) are computed for both sequence and structure similarity searches, making a total of six similarity search features. **B)** Pearson correlation coefficients (absolute values) between these six features and sequence identity in 100k human paralog pairs from Ensembl 111. **C)** Mean AUC of the six individual features compared to minimum sequence identity (in yellow) for predicting shared functions across the four datasets using four fold cross validation. Features with mean AUC greater than the mean AUC of sequence identity are shown in red. Error bars show the standard deviation of AUC values from cross-validation. **D)** Performances of an XGBoost classifier using the six similarity search features plus sequence identity (in green) compared to a classifier using solely sequence identity (in yellow), a classifier with only the three sequence similarity search features plus sequence identity (in purple), and a classifier with only the three structure similarity search features plus sequence identity (in light blue) for 4-fold cross-validation on shared functions prediction across the four datasets. Top chart shows the mean AUC (AUROC) on each dataset, bottom shows ROC and precision-recall curves for each datasets.

In human, Pearson correlation coefficients (Figure 5B) between these six features and sequence identity exhibited medium to low correlation with sequence identity (all < 0.5), indicating that these features successfully capture information not captured by sequence identity. Notably, no structure-based features exhibited correlation > 0.70 with sequence-based features, indicating that both similarity search approaches capture different information. Similar observations were made in yeast, as shown in Supplementary Fig. 1.

Since these six similarity search features capture sequence similarity of a paralog pair in the genomic and evolutionary context by relying on sequence/structure databases, we anticipated that they may also capture functional similarity. To evaluate their individual ability to predict if two paralogs share functions, we again performed 4-fold cross-validations on the four classification datasets. As shown in Figure 5C, most similarity search features had at least some predictive power (AUC > 0.5) and performed differently across all four datasets. Overall, the “paralog” features, those reflecting the presence of more similar sequences or structures to one of the two paralogs inside the genome, exhibited the strongest predictive power in most datasets.

Next, we combined all six similarity search features with minimum sequence identity in an XGBoost classifier and evaluated its performance using cross-validation on four datasets. As shown in Figure 5D, this similarity search classifier outperformed sequence identity alone in all datasets. The substantial improvement in all datasets when the 6 similarity search features are combined with sequence identity, compared to using sequence identity alone, highlights the presence of functional information captured by structure and sequence databases that pairwise sequence identity alone does not capture.

### Combining search and similarity features to better predict paralog pairs sharing functions

To capture the most comprehensive relationship between sequence divergence and functional divergence, we combined all previous features (structure similarity, PLM distances, and similarity searches) to model if two paralogs share functions in the two human datasets. Initially, we hypothesised that there might be a single latent variable underlying all the features that is predictive of shared functions and so we used the first dimension of a principal components analysis with all features to model the sequence divergence-functional divergence relationship. However, this approach performed poorly compared to most individual features and classifiers (Figure 6). This suggests that there is no single latent variable underlying all the similarity metrics, and we may need the full set of features to accurately capture ‘functional’ similarity. We then used all features to train machine learning classifiers (XGBoost) on the two human datasets. These classifiers achieved the best 4-fold cross-validation performances on the two human training datasets (Figure 6), significantly outperforming sequence identity alone or any of the classifiers trained on feature subsets (structural features, embedding features, database search features). This indicates that our features successfully capture different and unique aspects of sequence similarity between two protein sequences and their underlying functional relationships.

**Figure 6.**
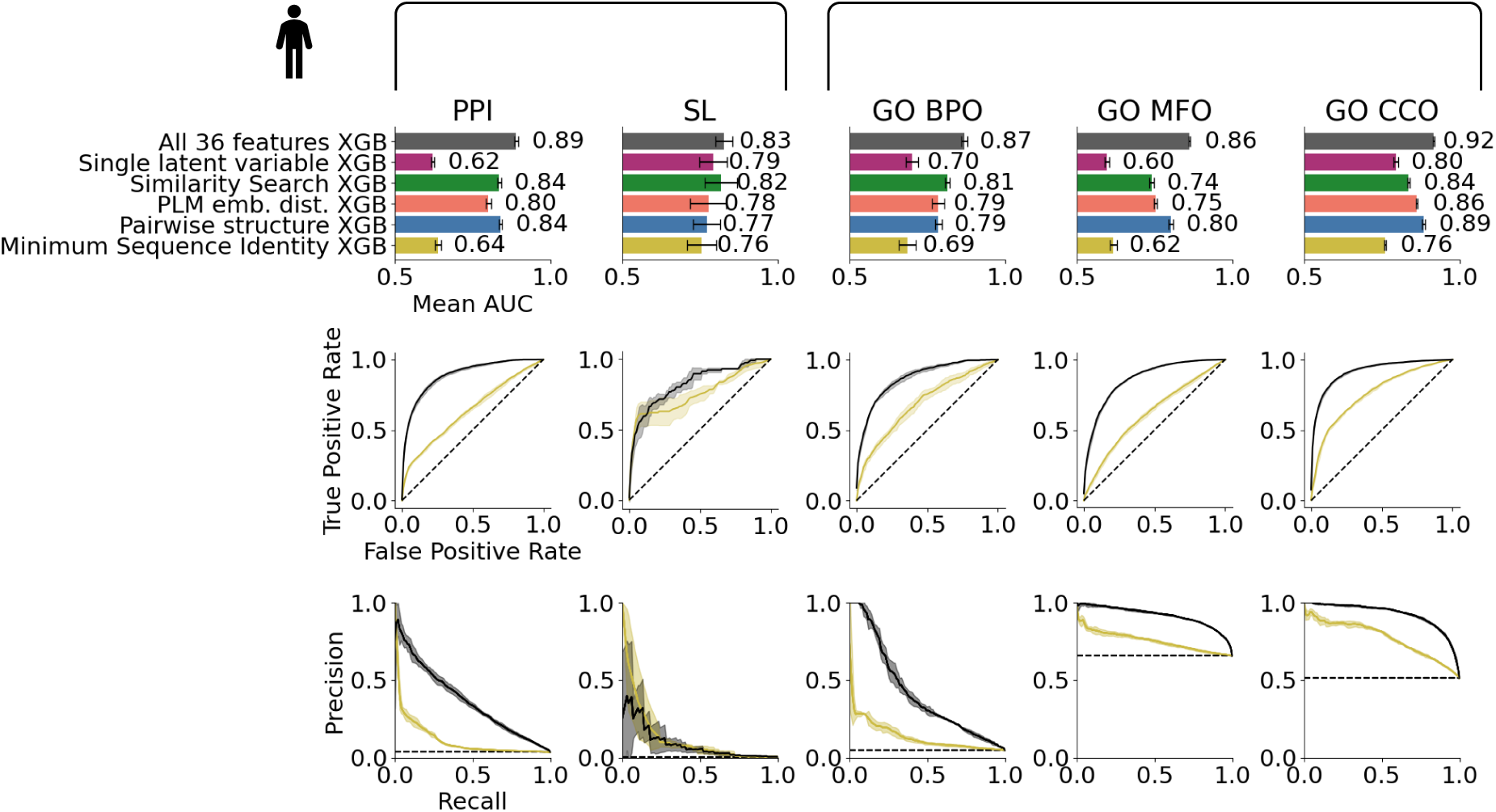
Integrating all features improves prediction of shared PPIs, synthetic lethality, and GO semantic similarity. Performances of an XGBoost classifier using all 36 sequence similarity features together (in grey/black) compared to a classifier using solely sequence identity (in yellow), the classifier using the nine predicted structure similarity features plus sequence identity (in blue), the classifier using the 20 PLM features plus sequence identity (in dark orange), the classifier using the six similarity search features plus sequence identity, and a classifier using only a single latent variable representing all 36 features (in purple) for 4-fold cross-validation on shared functions prediction across five human datasets (PPI ; SL ; GO BPO ; GO MFO ; GO CCO). Top chart shows the mean AUC (AUROC) on each dataset, bottom shows ROC and precision-recall curves for each datasets.

To validate these features and test if they can also capture different types of shared function definitions, we trained classifiers on three new human datasets based on the three Gene Ontology (GO) term categories: Biological Process (BPO), Molecular Function (MFO), and Cellular Component (CCO). We included only GO terms supported by experimental evidence to avoid class labels that were themselves based on sequence comparisons (See Methods). Paralog pairs were labelled as sharing functions (or not) based on the semantic similarity between their GO terms. In these three new tasks, our features significantly outperformed sequence identity alone (similar observations were made in yeast, as shown in Supplementary Fig. 2), validating them as invaluable sequence similarity metrics and demonstrating their ability to predict other definitions of shared paralog function.

## DISCUSSION

Recent AI advances in language processing have significantly impacted protein sequence analysis by introducing new representations, such as Protein Language Model (PLM) embeddings, and enabling accurate protein structure predictions, like those generated by AlphaFold2. In this study, we evaluated how well metrics derived from such models can predict shared functions in paralogs, compared to the traditional measure of sequence identity. Each of these features—whether comparing amino acid superposition in 3D structures, calculating distances between high-dimensional PLM embedding vectors, or assessing similarity through database searches—is based on different representations of protein sequences. Sequence identity compares the direct amino acid sequence, AlphaFold predicted structures capture co-evolutionary signals between amino acids, PLM embeddings represent sequences in a protein language space, and similarity searches compare sequences or structures against others in databases. Despite originating from the same protein sequences, we showed that these different metrics are non-redundant and they capture unique aspects of sequence divergence between paralogs. Furthermore, the combination of all these features in machine learning classifiers significantly outperforms any individual feature or feature type classifier across all datasets. While some studies may focus on specific types of features, such as structural data, PLM embeddings, or sequence-based approaches, our findings demonstrate that each type of feature captures different aspects of the sequence-function relationship. Our finding that the combination of all features performs well on tasks beyond shared PPI and SL prediction, i.e. predicting pairs with high Gene Ontology semantic similarity, suggests that the set of features we have calculated may be useful for other challenges that relate to paralog similarity, e.g. predicting common targets of small molecules (49). As a resource for the community, we therefore provide the features we have calculated for all paralog pairs on Zenodo at https://zenodo.org/records/13838296.

Both yeast and humans have undergone Whole Genome Duplication (WGD), yet they differ in the number of duplicated paralogs within each family. Human paralog families tend to be larger, potentially reflecting varying degrees of functional divergence. Despite these differences in duplication history, most features perform comparably in nearly all equivalent human and yeast datasets. The variations in feature performance may largely be attributed to differences in the dataset composition, with each dataset representing a distinct subset of the known paralog pairs. These subsets differ in their sequence identity distributions and the degree to which sequence and function are correlated. A key example is the human SL dataset constructed and studied in this paper. While it aims to be as comprehensive as current knowledge allows, human SL data remains less complete than that of yeast, where almost all paralog pairs have been systematically tested for SL. This means that some human SL paralog pairs may not be represented in the dataset, potentially limiting our ability to fully capture sequence/SL relationships, unlike in yeast. Nevertheless, across all datasets, the proposed sequence-derived features, especially in combination, effectively capture sequence-function relationships in both species. These promising results suggest the potential for broader application of these features beyond paralogs, such as predicting ortholog functions or determining whether two predicted orthologs share functions. Future work could explore this by applying our feature set to predict outcomes in protein replacement experiments (50–52), where one protein is substituted with its ortholog to observe functional conservation across species.

## DATA AVAILABILITY

*The data—including all computed features, datasets, and scripts used in this article—*are available on GitHub at https://github.com/cancergenetics/paralog_seq_similarity. *The produced features and datasets* are available on Zenodo at https://zenodo.org/records/13838296.

## ACKNOWLEDGEMENTS

We thank members of the Ryan lab for useful conversations and critical reading of the manuscript.

## FUNDING

This work was supported by a Science Foundation Ireland grant to CJR (20/FFP-P/8641). Funding for open access charge: Science Foundation Ireland.

## CONFLICT OF INTEREST

*Conflict of interest statement*. None declared.

## SUPPLEMENTARY DATA

**Supplementary Fig. 1.**
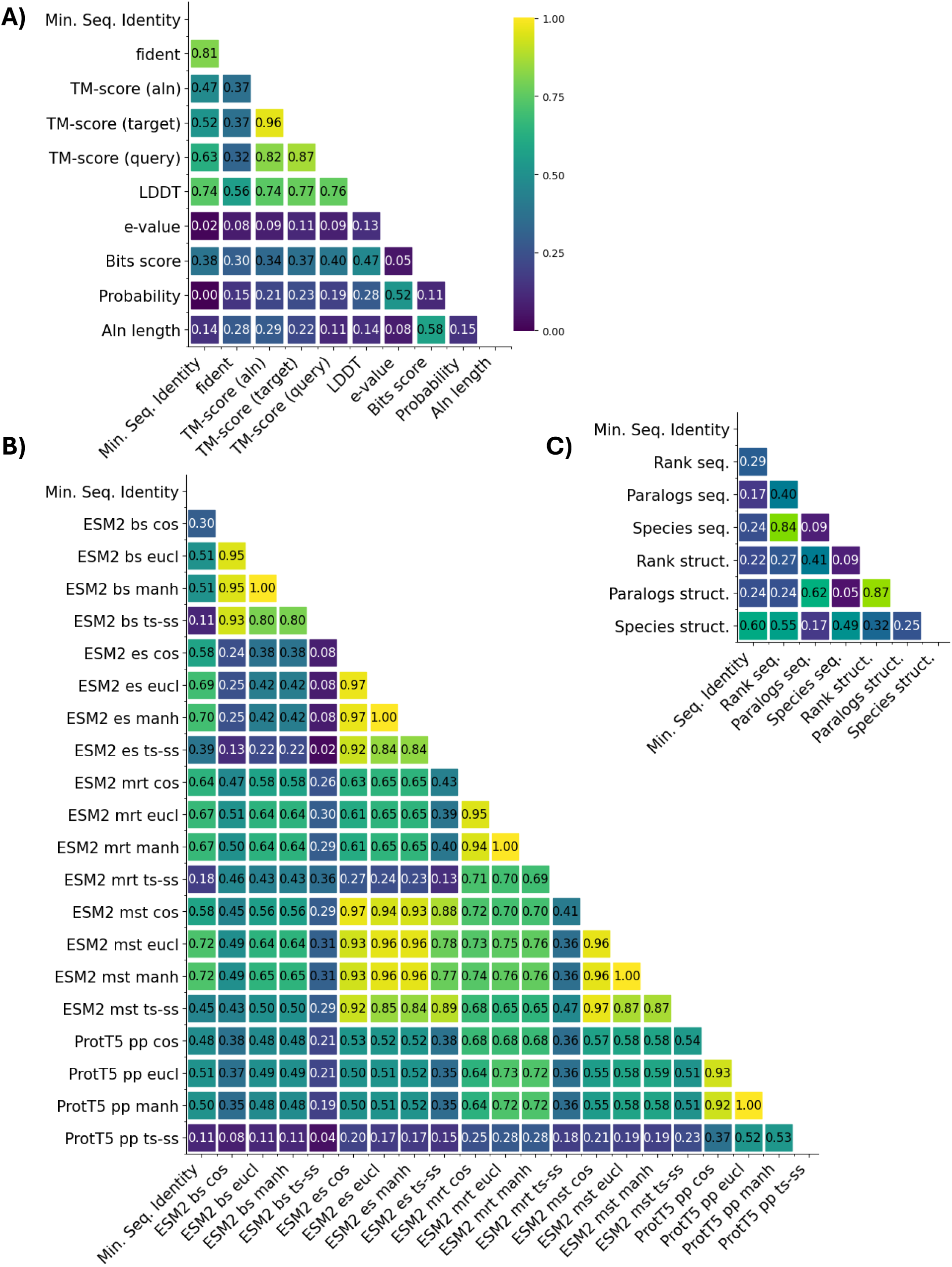
Pearson correlation coefficients (absolute values) between sequence features in 5,673 yeast paralog pairs. **A)** For the nine predicted structure similarity features and sequence identity. **B)** For the twenty Protein Language Model embedding features and sequence identity. **C)** For the six similarity search features and sequence identity.

**Supplementary Fig. 2.**
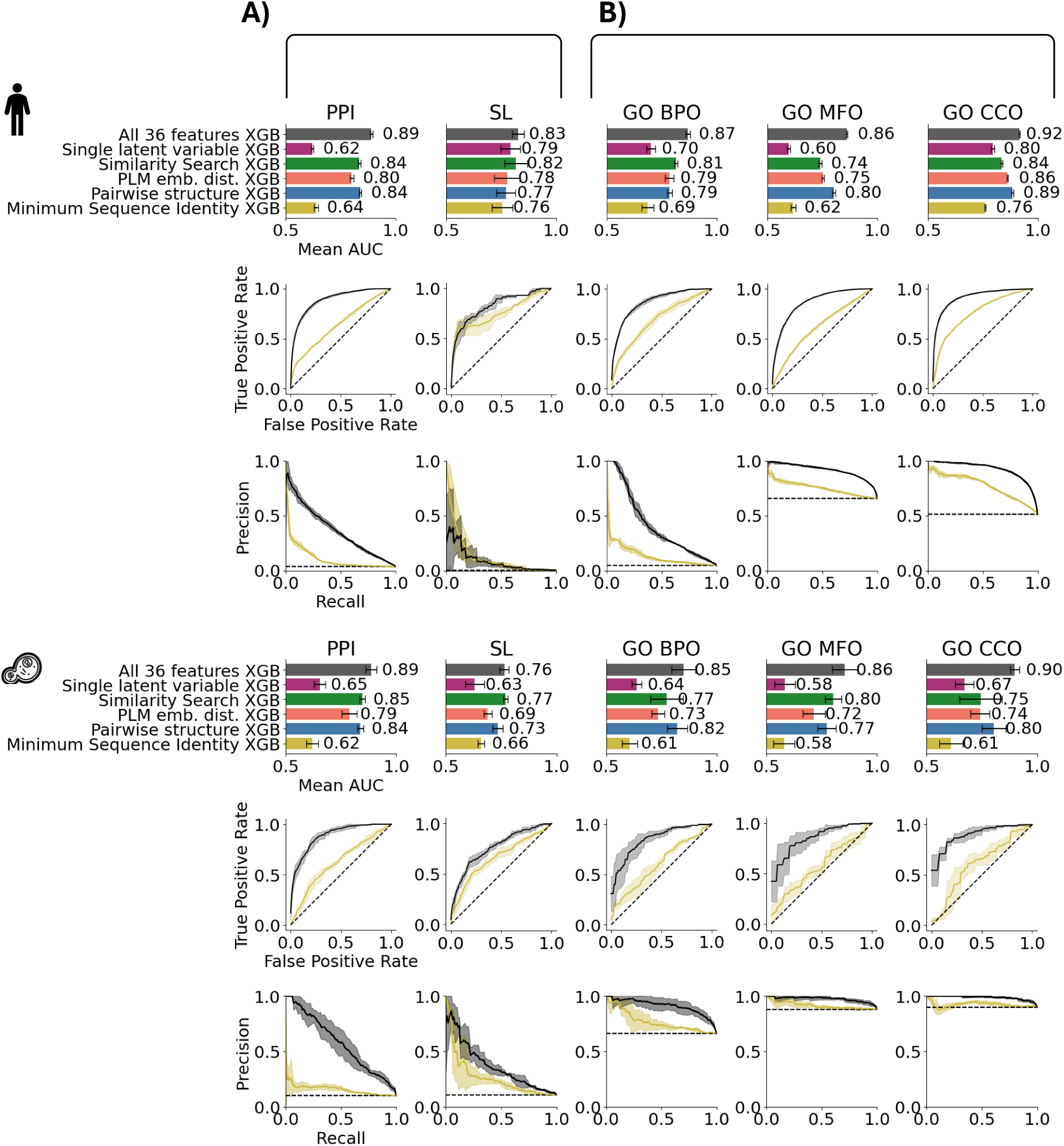
Integrating all features improves prediction of shared PPIs, synthetic lethality, and GO semantic similarity in human and in yeast. Performances of an XGBoost classifier using all 36 sequence similarity features together (in grey/black) compared to a classifier using solely sequence identity (in yellow), the classifier using the nine predicted structure similarity features plus sequence identity (in blue), the classifier using the 20 PLM features plus sequence identity (in dark orange), the classifier using the six similarity search features plus sequence identity, and a classifier using only a single latent variable representing all 36 features (in purple) for 4-fold cross-validation on shared functions prediction across five human datasets (PPI ; SL ; GO BPO ; GO MFO ; GO CCO) and their equivalent in yeast. Top chart shows the mean AUC on each dataset, ROC and precision-recall curves for each human datasets. Bottom chart shows the mean AUC on each dataset, ROC and precision-recall curves for each yeast datasets.

## Notes

https://zenodo.org/records/13838296

https://github.com/cancergenetics/paralog_seq_similarity

### Competing Interest Statement

The authors have declared no competing interest.

## REFERENCES

1. Altschul, S.F., Gish, W., Miller, W., Myers, E.W. and Lipman, D.J. (1990) Basic local alignment search tool. J. Mol. Biol., 215, 403–410.

2. Eisen, J.A. (1998) Phylogenomics: Improving Functional Predictions for Uncharacterized Genes by Evolutionary Analysis. Genome Res., 8, 163–167.

3. Eddy, S.R. (2009) A new generation of homology search tools based on probabilistic inference. Genome Inform. Int. Conf. Genome Inform., 23, 205–11.

4. Studer, R.A. and Robinson-Rechavi, M. (2009) How confident can we be that orthologs are similar, but paralogs differ? Trends Genet., 25, 210–216.

5. Nehrt, N.L., Clark, W.T., Radivojac, P. and Hahn, M.W. (2011) Testing the Ortholog Conjecture with Comparative Functional Genomic Data from Mammals. PLoS Comput. Biol., 7, e1002073.

6. Hernández-Salmerón, J.E. and Moreno-Hagelsieb, G. (2020) Progress in quickly finding orthologs as reciprocal best hits: comparing blast, last, diamond and MMseqs2. BMC Genom., 21, 741.

7. Ibn-Salem, J., Muro, E.M. and Andrade-Navarro, M.A. (2017) Co-regulation of paralog genes in the three-dimensional chromatin architecture. Nucleic Acids Res., 45, 81–91.

8. Dean, E.J., Davis, J.C., Davis, R.W. and Petrov, D.A. (2008) Pervasive and Persistent Redundancy among Duplicated Genes in Yeast. PLoS Genet., 4, e1000113.

9. Vavouri, T., Semple, J.I. and Lehner, B. (2008) Widespread conservation of genetic redundancy during a billion years of eukaryotic evolution. Trends Genet., 24, 485–488.

10. Iohannes, S.D. and Jackson, D. (2023) Tackling redundancy: genetic mechanisms underlying paralog compensation in plants. N. Phytol., 240, 1381–1389.

11. Ewen-Campen, B., Mohr, S.E., Hu, Y. and Perrimon, N. (2017) Accessing the Phenotype Gap: Enabling Systematic Investigation of Paralog Functional Complexity with CRISPR. Dev. Cell, 43, 6–9.

12. Hu, Y., Ewen-Campen, B., Comjean, A., Rodiger, J., Mohr, S.E. and Perrimon, N. (2022) Paralog Explorer: A resource for mining information about paralogs in common research organisms. Comput. Struct. Biotechnol. J., 20, 6570–6577.

13. Hartwell, L.H., Szankasi, P., Roberts, C.J., Murray, A.W. and Friend, S.H. (1997) Integrating Genetic Approaches into the Discovery of Anticancer Drugs. Science, 278, 1064–1068.

14. Kaelin, W.G. (2005) The Concept of Synthetic Lethality in the Context of Anticancer Therapy. Nat. Rev. Cancer, 5, 689–698.

15. DeLuna, A., Vetsigian, K., Shoresh, N., Hegreness, M., Colón-González, M., Chao, S. and Kishony, R. (2008) Exposing the fitness contribution of duplicated genes. Nat. Genet., 40, 676–681.

16. VanderSluis, B., Bellay, J., Musso, G., Costanzo, M., Papp, B., Vizeacoumar, F.J., Baryshnikova, A., Andrews, B., Boone, C. and Myers, C.L. (2010) Genetic interactions reveal the evolutionary trajectories of duplicate genes. Mol. Syst. Biol., 6, 429–429.

17. Han, K., Jeng, E.E., Hess, G.T., Morgens, D.W., Li, A. and Bassik, M.C. (2017) Synergistic drug combinations for cancer identified in a CRISPR screen for pairwise genetic interactions. Nat. Biotechnol., 35, 463–474.

18. O’Neil, N.J., Bailey, M.L. and Hieter, P. (2017) Synthetic lethality and cancer. Nat. Rev. Genet., 18, 613–623.

19. Lord, C.J. and Ashworth, A. (2017) PARP inhibitors: Synthetic lethality in the clinic. Science, 355, 1152–1158.

20. Kegel, B.D. and Ryan, C.J. (2019) Paralog buffering contributes to the variable essentiality of genes in cancer cell lines. PLoS Genet., 15, e1008466.

21. Dede, M., McLaughlin, M., Kim, E. and Hart, T. (2020) Multiplex enCas12a screens detect functional buffering among paralogs otherwise masked in monogenic Cas9 knockout screens. Genome Biol, 21, 262.

22. Huang, A., Garraway, L.A., Ashworth, A. and Weber, B. (2020) Synthetic lethality as an engine for cancer drug target discovery. Nat. Rev. Drug Discov., 19, 23–38.

23. Gu, Z., Steinmetz, L.M., Gu, X., Scharfe, C., Davis, R.W. and Li, W.-H. (2003) Role of duplicate genes in genetic robustness against null mutations. Nature, 421, 63–66.

24. Jumper, J., Evans, R., Pritzel, A., Green, T., Figurnov, M., Ronneberger, O., Tunyasuvunakool, K., Bates, R., Žídek, A., Potapenko, A., et al. (2021) Highly accurate protein structure prediction with AlphaFold. Nature, 596, 583–589.

25. Varadi, M., Anyango, S., Deshpande, M., Nair, S., Natassia, C., Yordanova, G., Yuan, D., Stroe, O., Wood, G., Laydon, A., et al. (2021) AlphaFold Protein Structure Database: massively expanding the structural coverage of protein-sequence space with high-accuracy models. Nucleic Acids Res., 50, D439–D444.

26. Rives, A., Meier, J., Sercu, T., Goyal, S., Lin, Z., Liu, J., Guo, D., Ott, M., Zitnick, C.L., Ma, J., et al. (2021) Biological structure and function emerge from scaling unsupervised learning to 250 million protein sequences. Proc. Natl. Acad. Sci., 118, e2016239118.

27. Elnaggar, A., Heinzinger, M., Dallago, C., Rehawi, G., Wang, Y., Jones, L., Gibbs, T., Feher, T., Angerer, C., Steinegger, M., et al. (2022) ProtTrans: Toward Understanding the Language of Life Through Self-Supervised Learning. IEEE Trans. Pattern Anal. Mach. Intell., 44, 7112– 7127.

28. Nallapareddy, V., Bordin, N., Sillitoe, I., Heinzinger, M., Littmann, M., Waman, V.P., Sen, N., Rost, B. and Orengo, C. (2023) CATHe: detection of remote homologues for CATH superfamilies using embeddings from protein language models. Bioinformatics, 39, btad029.

29. Zhao, C., Liu, T. and Wang, Z. (2024) PANDA-3D: protein function prediction based on AlphaFold models. NAR Genom. Bioinform., 6, lqae094.

30. Dickson, A. and Mofrad, M.R.K. (2024) Fine-tuning protein embeddings for functional similarity evaluation. Bioinformatics, 40, btae445.

31. Eisenberg, D., Marcotte, E.M., Xenarios, I. and Yeates, T.O. (2000) Protein function in the post-genomic era. Nature, 405, 823–826.

32. Harrison, P.W., Amode, M.R., Austine-Orimoloye, O., Azov, A.G., Barba, M., Barnes, I., Becker, A., Bennett, R., Berry, A., Bhai, J., et al. (2023) Ensembl 2024. Nucleic Acids Res., 52, D891–D899.

33. Huttlin, E.L., Ting, L., Bruckner, R.J., Gebreab, F., Gygi, M.P., Szpyt, J., Tam, S., Zarraga, G., Colby, G., Baltier, K., et al. (2015) The BioPlex Network: A Systematic Exploration of the Human Interactome. Cell, 162, 425–440.

34. Michaelis, A.C., Brunner, A.-D., Zwiebel, M., Meier, F., Strauss, M.T., Bludau, I. and Mann, M. (2023) The social and structural architecture of the yeast protein interactome. Nature, 624, 192–200.

35. Kegel, B.D., Quinn, N., Thompson, N.A., Adams, D.J. and Ryan, C.J. (2021) Comprehensive prediction of robust synthetic lethality between paralog pairs in cancer cell lines. Cell Syst, 12, 1144–1159.e6.

36. Costanzo, M., VanderSluis, B., Koch, E.N., Baryshnikova, A., Pons, C., Tan, G., Wang, W., Usaj, M., Hanchard, J., Lee, S.D., et al. (2016) A global genetic interaction network maps a wiring diagram of cellular function. Science, 353, aaf1420–aaf1420.

37. Varadi, M., Bertoni, D., Magana, P., Paramval, U., Pidruchna, I., Radhakrishnan, M., Tsenkov, M., Nair, S., Mirdita, M., Yeo, J., et al. (2023) AlphaFold Protein Structure Database in 2024:providing structure coverage for over 214 million protein sequences. Nucleic Acids Res., 52, D368–D375.

38. Kempen, M. van, Kim, S.S., Tumescheit, C., Mirdita, M., Lee, J., Gilchrist, C.L.M., Söding, J. and Steinegger, M. (2024) Fast and accurate protein structure search with Foldseek. Nat. Biotechnol., 42, 243–246.

39. Steinegger, M. and Söding, J. (2017) MMseqs2 enables sensitive protein sequence searching for the analysis of massive data sets. Nat. Biotechnol., 35, 1026–1028.

40. Consortium, T.U., Bateman, A., Martin, M.-J., Orchard, S., Magrane, M., Ahmad, S., Alpi, E., Bowler-Barnett, E.H., Britto, R., Bye-A-Jee, H., et al. (2022) UniProt: the Universal Protein Knowledgebase in 2023. Nucleic Acids Res., 51, D523–D531.

41. Heidarian, H. and Dinneen, D. (2016) A Hybrid Geometric Approach for Measuring Similarity Level Among Documents and Document Clustering. 2016 IEEE Second Int. Conf. Big Data Comput. Serv. Appl. (BigDataService), 10.1109/bigdataservice.2016.14.

42. Lin, Z., Akin, H., Rao, R., Hie, B., Zhu, Z., Lu, W., Smetanin, N., Verkuil, R., Kabeli, O., Shmueli, Y., et al. (2023) Evolutionary-scale prediction of atomic-level protein structure with a language model. Science, 379, 1123–1130.

43. Yeung, W., Zhou, Z., Mathew, L., Gravel, N., Taujale, R., O’Boyle, B., Salcedo, M., Venkat, A., Lanzilotta, W., Li, S., et al. (2023) Tree visualizations of protein sequence embedding space enable improved functional clustering of diverse protein superfamilies. Brief. Bioinform., 24, bbac619.

44. Cherry, J.M., Hong, E.L., Amundsen, C., Balakrishnan, R., Binkley, G., Chan, E.T., Christie, K.R., Costanzo, M.C., Dwight, S.S., Engel, S.R., et al. (2012) Saccharomyces Genome Database: the genomics resource of budding yeast. Nucleic Acids Res., 40, D700–D705.

45. Zhao, C. and Wang, Z. (2018) GOGO: An improved algorithm to measure the semantic similarity between gene ontology terms. Sci. Rep., 8, 15107.

46. Braun, P., Tasan, M., Dreze, M., Barrios-Rodiles, M., Lemmens, I., Yu, H., Sahalie, J.M., Murray, R.R., Roncari, L., Smet, A.-S. de, et al. (2009) An experimentally derived confidence score for binary protein-protein interactions. Nat. Methods, 6, 91–97.

47. Zhang, Y. and Skolnick, J. (2004) Scoring function for automated assessment of protein structure template quality. *Proteins: Struct., Funct.*, Bioinform., 57, 702–710.

48. Mariani, V., Biasini, M., Barbato, A. and Schwede, T. (2013) lDDT: a local superposition-free score for comparing protein structures and models using distance difference tests. Bioinformatics, 29, 2722–2728.

49. Kruger, F.A. and Overington, J.P. (2012) Global Analysis of Small Molecule Binding to Related Protein Targets. PLoS Comput. Biol., 8, e1002333.

50. Kachroo, A.H., Laurent, J.M., Yellman, C.M., Meyer, A.G., Wilke, C.O. and Marcotte, E.M. (2015) Systematic humanization of yeast genes reveals conserved functions and genetic modularity. Science, 348, 921–925.

51. Laurent, J.M., Garge, R.K., Teufel, A.I., Wilke, C.O., Kachroo, A.H. and Marcotte, E.M. (2020) Humanization of yeast genes with multiple human orthologs reveals functional divergence between paralogs. PLoS Biol., 18, e3000627.

52. Lai, H.-Y., Yu, Y.-H., Jhou, Y.-T., Liao, C.-W. and Leu, J.-Y. (2023) Multiple intermolecular interactions facilitate rapid evolution of essential genes. *Nat*. Ecol. Evol., 7, 745–755.

