## Supplementary figures and images for "Evaluating Sequence and Structural Similarity Metrics for Predicting Shared Paralog Functions"

### Supplementary Fig. 1

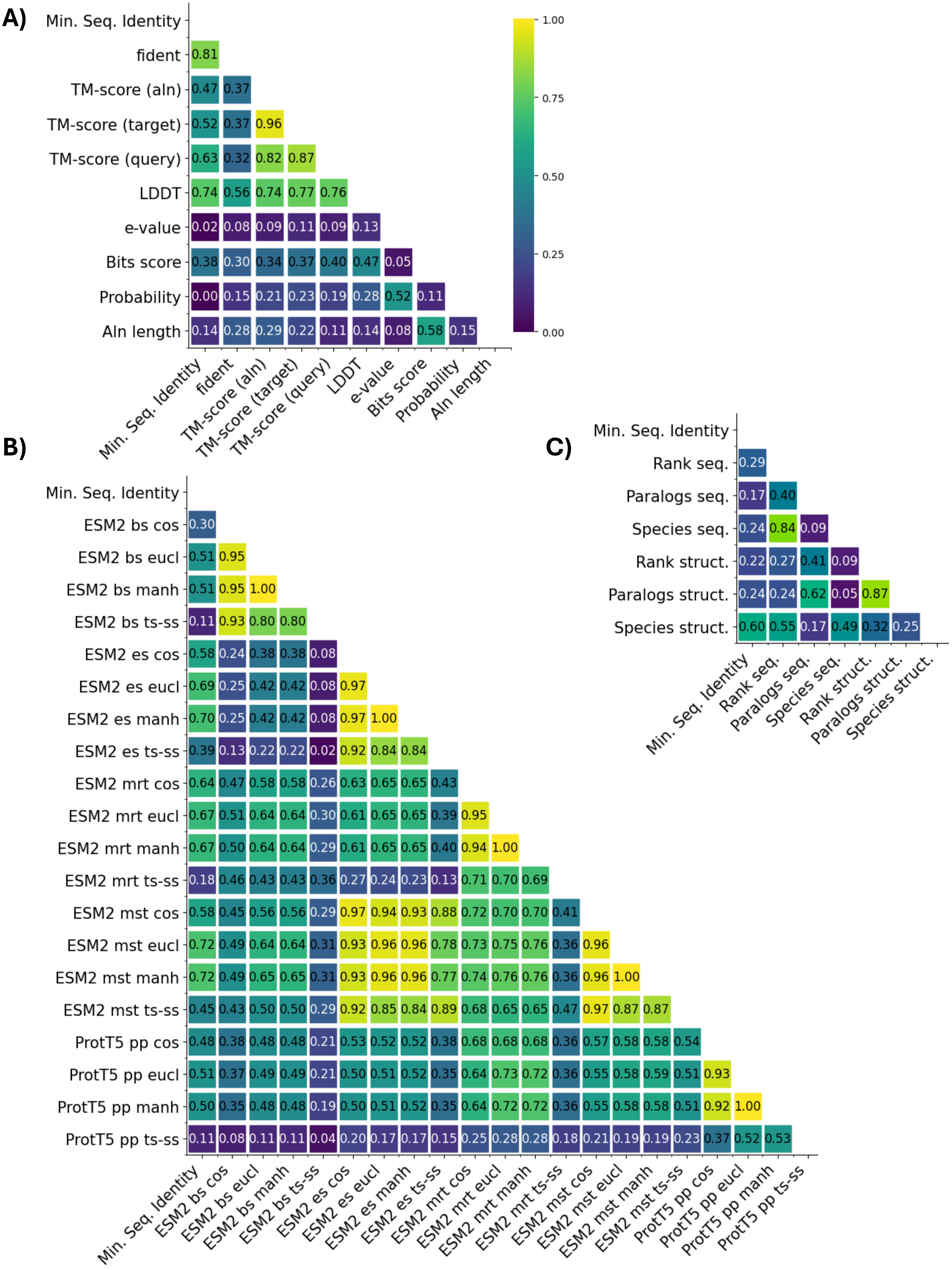

### Supplementary Fig. 2

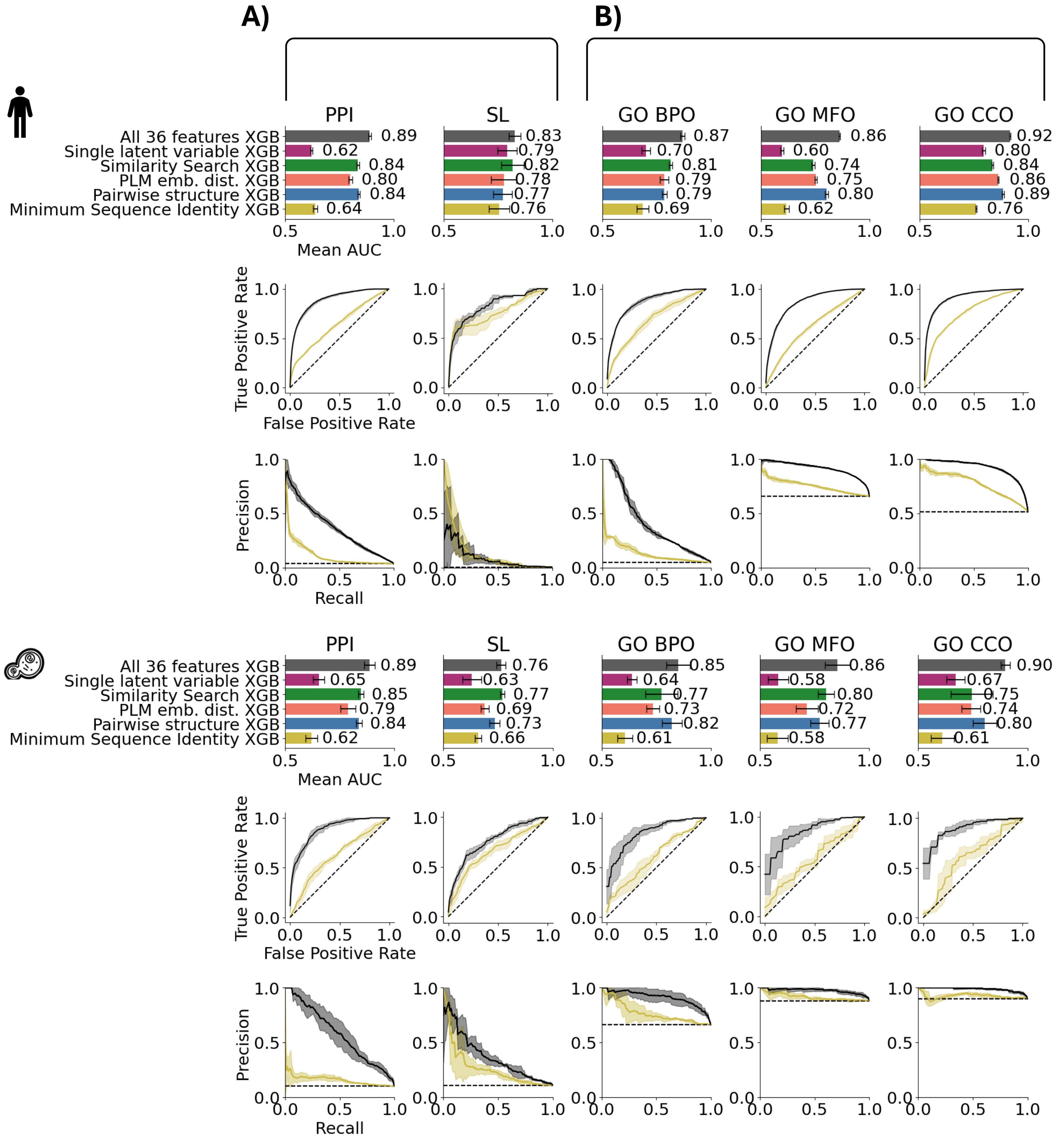
